# Adaptation of the Geometric Viability Assay as a high throughput method to measure *Clostridioides difficile* spore viability and germinant sensitivity

**DOI:** 10.64898/2026.10.19.760248

**Authors:** Kyle Vogtman, Amy Rohlfing

## Abstract

*Clostridioides difficile* is the leading cause of hospital acquired infections and gastroenteritis associated death in the United States. As an obligate anaerobe, *C. difficile* relies on the metabolically dormant, aerotolerant spore form to transmit infection. *C. difficile* infection begins when spores germinate in the small intestine in response to specific bile acids and a co-germinant signal (either amino acids or divalent cations). As germination is required to initiate infection, understanding how spores sense germinants and outgrow into toxin-producing cells is an active area of research. One of the fundamental assays used to study *C. difficile* germination is a resource and time intensive assay, similar to a bacterial CFU assay, that relies on serial dilutions of a spore sample plated on agar media in triplicate to determine the number of viable spores in a sample. As this assay is similar to assays used to enumerate viable bacteria in a sample, we hypothesized that a recently developed high-throughput technique to measure bacterial CFUs published by Meyers *et al.,* the Geometric viability assay (GVA), could be modified to measure the number of viable spores in a sample. We compared measurements of viable spores per mL from the conventional spore viability assay and the modified GVA protocol and found that the two measurements were highly consistent with each other (Pearson r=0.991, R^2^=0.9821). The modified GVA protocol was also able to measure differences in sensitivity to germination signals in strains of *C. difficile* hypersensitive to co-germinants and measure low levels of spontaneous germination. The GVA protocol is less resource and time intensive than the conventional assay, was easy adapt, and likely would be feasible to study germination in a wide range of spore-formers.

## Introduction

*Clostridioides difficile* is a Gram-positive, spore-forming, obligate anaerobe(1). Classified by the CDC as an urgent antibiotic threat, *C. difficile* is estimated to have caused 223,900 hospitalized infections and 12,800 deaths in 2017(2–4). *C. difficile* is the leading cause of hospital acquired infections in the United States, contributing to over 15% of all hospital acquired infections and 22% of those with an associated pathogen (5). While most individuals are protected from *C. difficile* infection by the diverse microflora in the gastrointestinal tract, hospitalized individuals receiving broad-spectrum antibiotics can become susceptible to *C. difficile* infection due to disruption of the gut microbiome (1, 6).

As *C. difficile* is an obligate anaerobe, the aerotolerant and resistant spores are required to transmit infection between individuals (7). *C. difficile* spores also contribute to recurrent infections. *C. difficile* spores are inert to antibiotics and can outlast antibiotic treatment to resolve a *C. difficile* infection, seeding a future infection (8–11). *C. difficile* infection begins when the spores detect specific environmental signals in the small intestine of a susceptible individual, triggering the process of germination (12–14). Germinating spores undergo large morphological changes to outgrow and enter the vegetative state (12, 15, 16). *C. difficile* vegetative cells are then able to colonize the colon and produce inflammatory toxins, which are the primary virulence factors (17, 18). As infection progresses, unknown factors trigger sporulation (1). When the spores exit the host, they can initiate the infection cycle in another susceptible individual or, in about 20% of cases, infect the same individual to cause a recurrent infection (3). Thus, because of the anaerobic nature of *C. difficile*, germination is critical to the infection cycle (9–11).

*C. difficile* has a unique germination signaling cascade. While most spore forming bacteria germinate in response to nutrient signals, *C. difficile* germinates in response to specific mammalian bile acids, particularly taurocholic acid (TA) (19, 20). In addition, *C. difficile* spores require a second co-germinant signal, either amino acids or divalent cations, to germinate (20, 21). How *C. difficile* spores sense and respond to these germination signals is also unique. C*. difficile* is one of only two known spore formers that lacks membrane bound Ger-type receptors to sense germination signals (22). Instead, the *C. difficile* spore germination signaling cascade consists of three subtilisin-like serine proteases, CspA, CspB, and CspC (13, 19, 23–25). While the Csp family of proteases is conserved in other Clostridia, in *C. difficile*, only CspB is an active protease (26, 27). The CspA and CspC encoded in *C. difficile* have mutations in the conserved catalytic triad, rendering them inactive pseudoproteases (23, 25). Recently published work found that CspA and CspC form a stable heterodimer and determined that the CspA-CspC heterodimer is likely the active form that senses bile acid germinants and propagates the signal (28). Indeed, another recent study published a Cyro-EM structure of the CspA-CspC heterodimer with the bile acid taurocholate bound at three sites within the CspA-CspC heterodimer, with two of these sites along the interface of the heterodimer (29).

After sensing germinants, the CspA-CspC heterodimer activates the protease activity of CspB through an unknown mechanism (23, 26). CspB is then able to proteolytically activate the cortex hydrolase, SleC, to begin the large morphological changes that occur during germination, including degradation of the thick cortex, release of the spore-specific molecule calcium dipicolinic acid (CaDPA) from the spore core, and rehydration of the spore core (12, 23, 26, 30–32). Despite these recent advances in our understanding of the molecular mechanisms controlling germination initiation, many questions remain regarding germinant sensing and the signaling cascade that eventually leads to resumption of metabolic activity. A thorough understanding of these processes could aid the development of novel therapeutics that specifically target and inhibit the unique germination pathway of *C. difficile* (14). Targeted therapies against *C. difficile* spore germination are proposed to reduce further disruption to the gut microbiome and limit recurrent infections (14, 33).

One factor limiting research into understanding the complete process of spore germination from germinant sensing to resumption of metabolic activity is the lack of a high-throughput method to measure spore germination and outgrowth. There are three major techniques used in studies of *C. difficile* spore germination: an optical density (OD)-based assay that measures cortex degradation and core rehydration (20, 34), a fluorescence-based assay to measure release of CaDPA after cortex degradation (31, 35), and a spore viability assay similar to a colony-forming unit (CFU) spot assay (34). While the OD-based and fluorescence based assays can be completed in 96-well plates to be scaled up for high-throughput studies, these assays rely on bulk population measurements and only measure early germination events (35–38). The OD-based assay to measure cortex degradation and core hydration in particular misses germination events that occur at lower frequency (37). On the other hand, the spore viability assay measures the concentration of spores that are able to both germinate and outgrow into viable colonies and can capture germination events that happen at a lower frequency (23, 34).

The spore viability assay involves serially diluting a sample of spores and plating a small volume of the dilutions onto agar medium containing the bile acid germinant taurocholate (34). The spots are allowed to grow overnight, and colonies are counted to determine the concentration of viable spores in the solution. In addition to assessing the number of viable spores in a sample, this assay can also be modified to determine the sensitivity of spores to germination signals (24, 39). However, the spore viability assay has many of the same limitations as the spot CFU assay. Primarily, these methods are resource- and time-intensive. Each assay requires numerous pipette tips to complete the serial dilutions and an agar media plate for each sample or condition tested (40). These material costs quickly add up over a large number of assays. Furthermore, the necessary serial dilutions are time-consuming and a point where error can be introduced. This limits the feasibility of large-scale experiments and makes high-throughput experiments difficult if not impractical (40).

The Geometric Viability Assay (GVA) recently developed by Meyer, *et al.* is a high-throughput assay that can replace the CFU spot assay to enumerate viable bacteria within in a sample (40). As the spore viability assay procedure is similar to that of the CFU spot assay and the two procedures share limitations, we hypothesized that GVA could be adapted to measure the number of viable *C. difficile* spores in a sample. GVA removes the need to generate serial dilutions to create a concentration gradient of cells for enumeration. Instead GVA utilizes the probability gradient that naturally occurs within a cone, such as a pipette tip, in order to calculate the number of CFUs within a sample. The probability gradient naturally forms in a cone due to the constant decrease in cross sectional area as one moves further from the base of the cone(40). Meyer et al. utilized the probability encoded in the position in which colonies grew in a pipette tip to calculate the number of CFUs in a sample (40). GVA method was able to reduce both the materials and time required to calculate the CFUs in a solution, with a reported 36x reduction in time during the plating step and 13x reduction in total cost as compared to a CFU spot assay (40). Other work with this technique has shown that the same principle can be applied to anaerobic bacteria (41).

In this study, we adapt GVA method to measure the viability of *C. difficile* spores. We found that measurements the number of viable spores in a sample was comparable between the spore viability assay and GVA and that GVA was accurate across 5 orders of magnitude. We found that while the upper limit of detection of GVA was lower than that of the spore viability assay, GVA assay had a lower limit of detection and could measure viable spores down to around 26 CFUs mL^-1^. We were also able to adapt GVA to measure differences in sensitivity to germinants between *C. difficile* strains and capture low frequency spontaneous germination events that occur in the absence of bile acid germinants. In modifying the GVA optical configuration for our laboratory, we were also able to increase the flexibility of the optical set-up for use with a wider variety of macro lens, smart phones and pipette tips. Overall, GVA is a time- and resource-saving assay for accurately measuring *C. difficile* spore viability, spore germinant sensitivity, and low-level germination events.

## Results

### Optimization an Optical Configuration for use of GVA with *C. difficile*

Quantification of the Geometric Viability Assay (GVA) requires an optical setup to capture images of colonies grown overnight in media embedded in a pipette tip. There are four methods for imaging pipette tips that have been developed for use with GVA: 1) a digital mirrorless camera with a 1:1 macrolens and motorized stage, 2) a smart phone with clip-on microlens and 3D printed holder for phone and pipette tip, 3) a printed ruler and magnifier, or 4) a GelDoc imager (40, 41). Based on the availability of 3D printing to the authors and the low cost of a clip-on smart phone macro lens, the smartphone optical configuration designed by Meyer *et al.* was selected as the base for our optical configuration design (40). While the clip-on macrolens and smart phone optical setup produce lower resolution images than the mirrorless camera and macrolens option, Meyer *et al.* determined that the accuracy of GVA is not impacted by the imaging method, although the smartphone and clip-on macrolens configuration does have a lower upper limit of detection compared to the digital camera setup.

The 3D printed phone and tip holder originally described by Meyer *et al*., had three main components: 1) a base to hold a smart phone, 2) a component that fit snugly around the clip-on macrolens that the pipette tip holder attaches to, and 3) a pipette tip holder that could be adjusted both vertically and horizontally to place the pipette tip in frame of the smart phone camera (40). In testing this original design, we had difficulties securing the pipette tip holder attachment device (component 2), as slight differences in how it was printed resulted in too loose a fit. This resulted in the attachment rotating to incorrect angles and the pipette tips not being perpendicular to the ground. This design also limited the macro lenses that could be used to the one used in the original manuscript. Additionally, the hole that held the pipette tip in the pipette tip holder (component 3) had a diameter slightly larger than optimal. This resulted in the pipette tips shaking when placed in the holder and caused blurry photos. To resolve the first issue and make the setup usable with any macro lens, including built-in macro lenses on a smartphone, we modified the design of the 3D printed holder to attach the height adjustor/tip holder directly to the base of the optical configuration (Fig. 1A). This gave the configuration greater flexibility with a combination of smart phones and macrolenses and increased stability and consistency while imaging. We also modified the diameter of the pipette tip holder to better fit the brand of pipette tips (TipOne from USA Scientific) used by our laboratory (Fig. 1B). These modifications resulted in clearer photos and faster imaging, as fewer adjustments were needed between pipette tips. All experiments described in this manuscript were performed using the modified set-up with the tip holder attached to the base.

**Figure 1:**
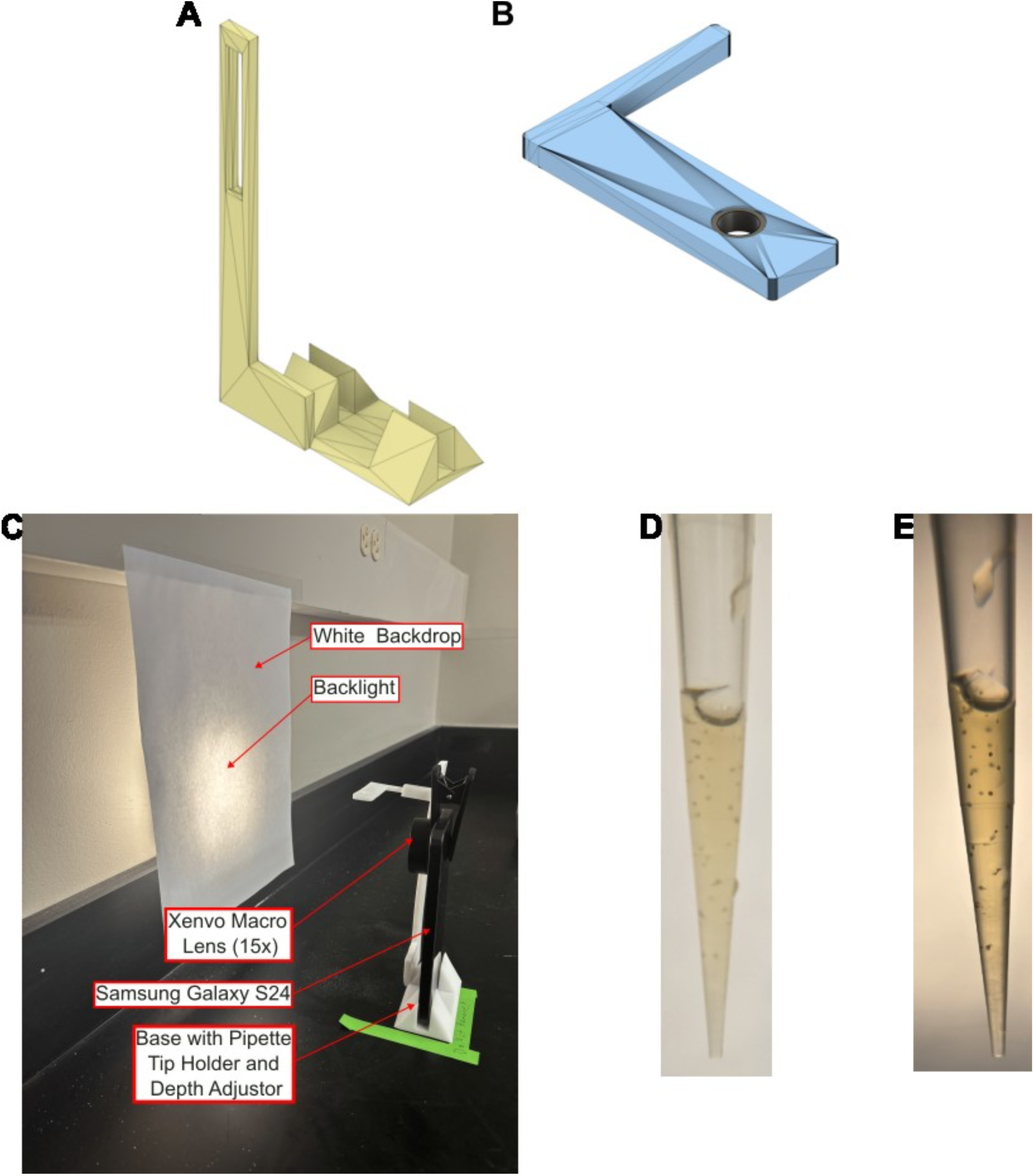
Modifications were made to the original optical configuration to improve image quality, replicatbility, and image contrast **(A)** The base of the optical configuration was modified to the height adjustor (vertical position) to the phone holding portion. This allows for any phone and macro lens combination to be used. **(B)** The hole in the pipette tip holder can be modified to better fit any brand of pipette tip, such as TipOne, the optimized brand for the holder displayed. **(C)** Fully assembled optical configuration. Using a back light in addition to the ambient light improved colony contrast. **(D)** Without the backlight, the colonies have low contrast against the background. **(E)** Adding the backlight increased the contrast of the colonies in the same pipette tip and expands the dynamic range by one order of magnitude.

Initial images with the modified 3D printed image configuration revealed low contrast between colonies and the image background (Fig. 1D). The low contrast of the images was particularly detrimental when imaging tips with high CFU counts as these tips had small, individual colonies at the end of the pipette tips that were difficult to resolve. In initial tests, we were only able to quantify spore samples within four orders of magnitude, one order of magnitude lower than that of the smartphone configuration created by Meyer et al (40). To increase contrast of colonies grown anaerobically, El-Fateh, *et. al* had added the pH sensitive dye bromocresol green to the media (41). However, we found that *C. difficile* growth did not result in a large enough pH change of the supplemented BHI media for a color change in the bromocresol green to be noted. We next tested a physical method of increasing contrast by adding a backlight to the imaging set-up. A small rechargeable LED light was positioned behind the white paper used as a backdrop. This simple solution was able to increase the contrast of the colonies compared to the background (Fig. 1E). Figures 1D and 1E display this difference with a picture of the same tip with and without the backlight. While the increased colony contrast increased the ability of the GVA software to properly identify colonies, colonies towards the outside “edge” of the pipette tip were often obscured by dark shadows. However, we hypothesized that our imaging set-up would still produce accurate results as Meyer et al. reported that 10 of 15 colonies could be missed and the result would not be statistically different from if all the colonies had been counted (40). Thus, we continued with the final optical configuration (Fig. 1C) to test the ability of the GVA to measure viability and germinant sensitivity of *C, difficile* spores.

### Measuring the number of viable spores in a sample using the GVA assay

We first tested if the GVA could measure differing concentrations of spores in a sample by comparing the measurements of samples of varying concentrations of spores to the measurements of the spore viability assay. To do this, *C. difficile* 630Δ*erm* spores were diluted to a concentration of approximately 1×10^7^ spores mL^-1^ (0.35 OD_600_ units in 1 mL) to generate a solution with a known concentration of spores. This spore solution was then serially diluted 11 times by a factor of 5, down to approximately 2×10^-1^ spores mL^-1^. Each of these dilutions was treated as a sample. The concentration of spores in each “sample” was then measured in biological and technical triplicate using both a spore viability assay and GVA. The spore viability assay was performed as previously described (32). Briefly, each spore sample was serially diluted 10-fold six times. The six 10-fold dilutions were plated in triplicate on supplemented BHI media containing a 0.1% (1.9 mM) taurocholate germinant (BHIS+TA). Spot plates were incubated for about 22 hours at 37°C anaerobically. Colonies were counted in dilutions with 10-25 colonies to calculate the number of viable spores in the sample.

GVA was performed similarly to the originally described method (40), with BHI medium supplemented with yeast extract (5g/L), L-cystine (0.1% w/v) (BHIS) and the germinant taurocholate (0.1% w/v) with 0.66% agarose concentration as the growth media for embedding. The medium was autoclaved and allowed to solidify. Before use, the agarose-containing medium was microwaved until liquid and then allowed to cool to 55°C in a water bath in an anaerobic environment during embedding. 150µL of molten medium was mixed with 50 µL of diluted spore sample, resulting in a 1:4 dilution; 150µL of sample-media mixture was then aspirated into a 200µL pipette tip. The tip was submerged in an ice bath for approximately 20 seconds and then allowed to sit at ambient temperature in the anaerobic chamber for approximately 30 minutes. Following this, the tips were placed in an anaerobic incubator and allowed to grow overnight. After growth, the tips were removed from the anaerobic chamber and imaged using a Samsung Galaxy S24 with a Xenvo Macro Lens. Following imaging, the photos were analyzed with the GVA software (40) and CFUs mL^-1^ were calculated. When paired with a spore viability assay, the tips were placed in and removed from the incubator at the same time.

The GVA measurement of each sample was plotted against the spore viability measurement for the same sample (Fig. 2A) and a line of best fit was determined through linear regression. For samples where GVA had visible colonies, but the spore viability assay had none, the spore viability assay value was set to the last readable spore viability reading divided by 5 to the power of the number of dilutions after that last recorded measurement. This gave an approximation of the number of spores that should be found in sample for statistical analysis. A Pearson test and Bland-Altman analysis were also performed in order to compare the results. One outlier indicated in Fig. 2A was excluded from statistical analysis and the linear regression. This result was excluded from analysis as it was well outside of the 95% confidence interval on the Bland-Altman analysis and had a value at concentrations below the lower limit of detection. Linear regression found a line of best fit with a slope of 0.96. This was found to be statistically indistinguishable from one, the expected slope (t test, p = 0.16). The Pearson test also confirmed high correlation between the measurements (Pearson r = 0.99, p = 1×10^-20^). This high correlation also indicates that the hot media had no statistical effect on the viability of the spores. A Bland-Altman analysis (Fig. 2B) was performed and revealed an average bias in GVA of 10^0.22^(1.7) CFUs mL^-1^, indicating that GVA measured an average of 10^0.22^ CFUs/mL more on average than the spore viability assay. The trendline of the difference was shown to be statistically indistinguishable from zero (t test, p=0.28), a characteristic of homoscedastic data.

**Figure 2:**
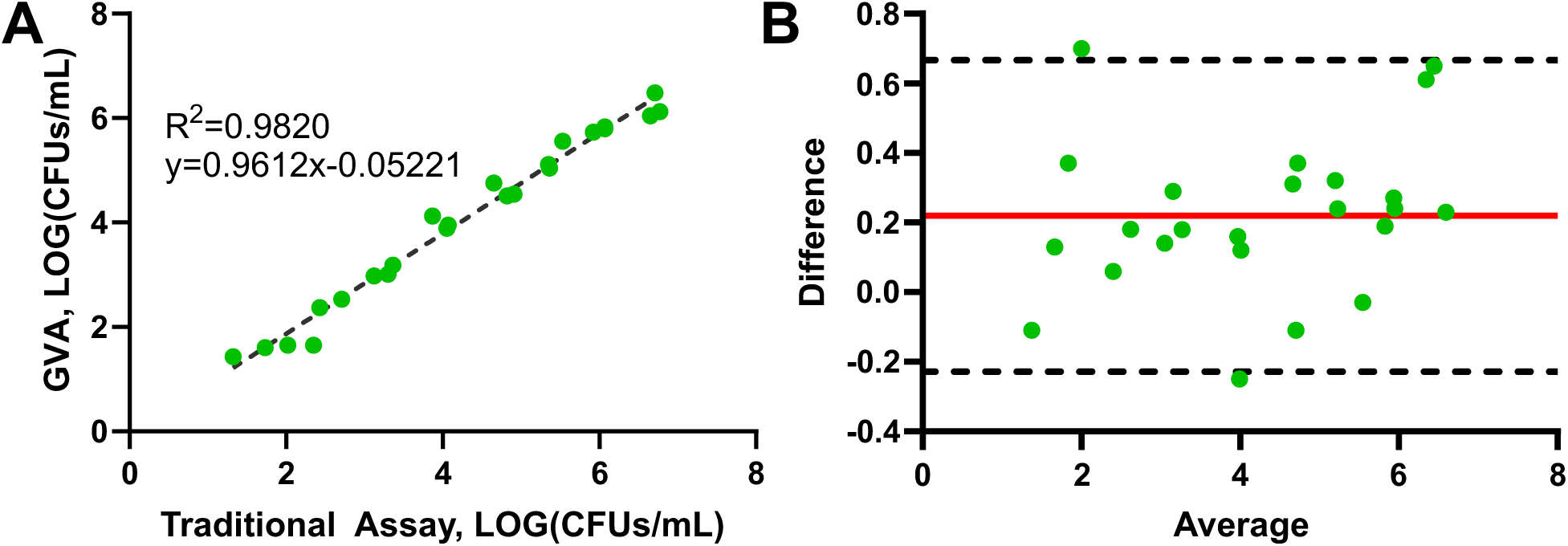
GVA and traditional assay measurements were compared using linear and Bland-Altman analysis. **(A)** The spore viability assay and GVA measurements are highly correlated across five orders of magnitude. Each datapoint represents the average of three technical replicates. Dashed line is the line of best fit Correlation was confirmed across three biological replicates. One outlier was noted (not presented) and was excluded as the traditional assay measurement was well below the LOD of the GVA (Traditional Assay: -0.08, GVA:1.73 LOG(CFU/mL)). **(B)** Bland-Altman analysis was used to calculate the average bias of the GVA, represented with a red line, which was found to be 10^0.22^, or 1.7 CFUs/mL. The dashed line indicates the 95% confidence interval. The slope of the trendline for the data was found to be statistically indistinguishable from zero (T-test, P = 0.28) indicating homoscedastic data with minimal differences.

From these results, we also determined the dynamic range of the modified GVA protocol (Fig. 3). The images of the pipette tips had individually distinguishable colonies for concentrations of spores up to approximately 1×10^6^ spores mL^-1^. The reported lower limit of detection for the specifications we used was 26 CFUs mL^-1^(40). The dynamic range determined from this spanned 5 orders of magnitude, the same range as Meyer et al. were able to achieve with their smart phone optical setup(42). This dynamic range is smaller than the dynamic range of the spore viability assay but is able to measure two orders of magnitude lower CFU mL^-1^ than the spore viability assay.

**Figure 3:**
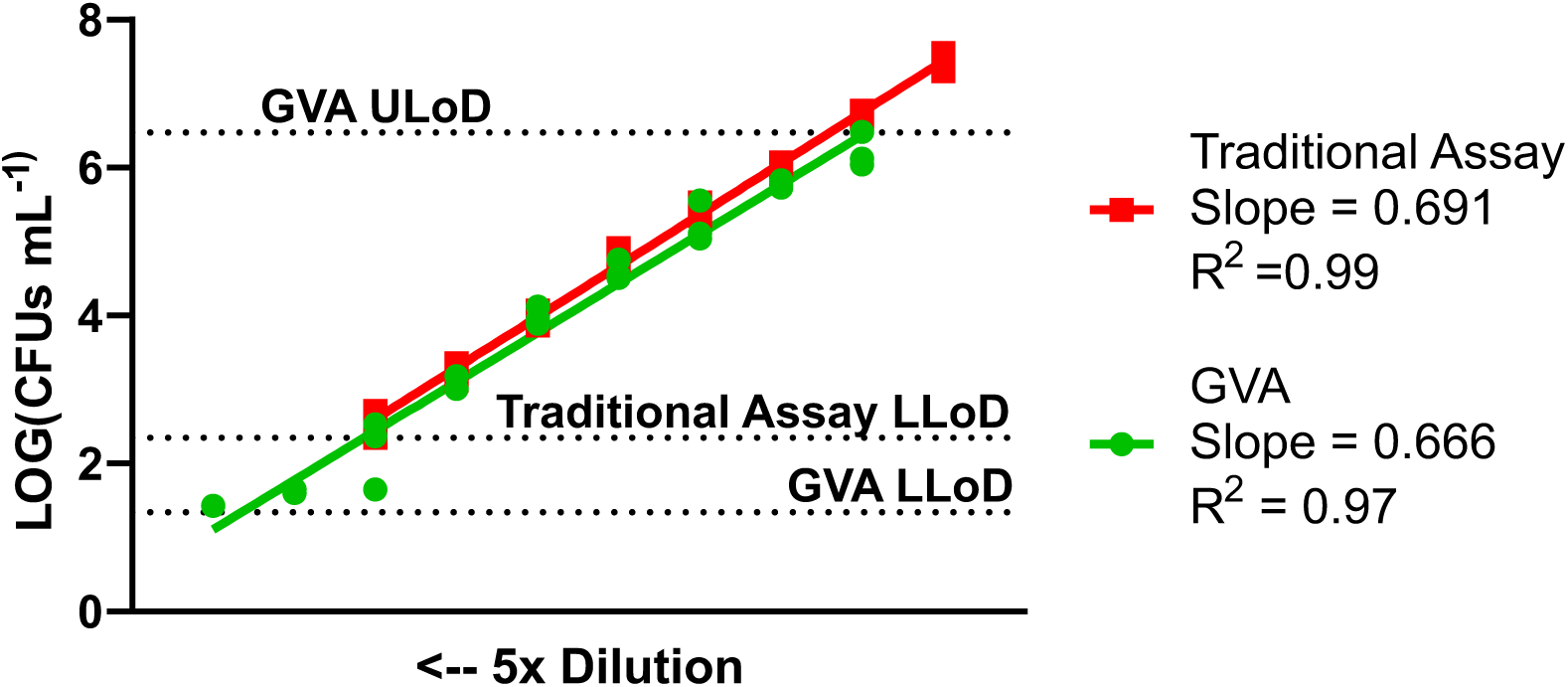
The lower limits of detection (LLoD) of GVA (22 CFUs mL^-1^) and the traditional spore viability assay (222 CFUs mL^-1^) differ by approximately one order of magnitude. The dynamic ranges of each assay were found by generating a series of five-fold dilutions (10 total dilutions enumerated with viable colonies). The dynamic range of GVA spans five orders of magnitude (ULoD approximately 3.0 x 10^6^ CFUs mL^-1^) while the spore viability assay spans 7 orders of magnitude (theoretical ULoD at 2.2 x 10^9^ CFUs mL^-1^).

### The GVA assay measures rare spontaneous germination events

As our initial testing of the GVA to measure viability of *C. difficile* spores indicated that GVA had a lower limit of detection than the spot dilution spore viability assay, we wanted to test the ability of GVA to measure the low rate of spontaneous germination (or germination without bile salt germinant) of *C. difficile* spores that occurs on agar containing media. We tested three previously described strains of *C. difficile* that differ in their response to germinants and/or spontaneous germination rates to determine if GVA would accurately measure these differences. We compared the number of spores that spontaneously germinate between 630Δ*erm* spores, spores with a deletion of the gene for the bile acid germinant receptor, *cspC* (ΔcspC), and spores that encode a CspC variant that is hypersensitive to co-germinant signals (ΔcspC/cspC(G457R)) (19, 24). Δ*cspC* spores are unable to respond to bile acid germinants (19) and thus are only able to germinate spontaneously (23). ΔcspC/cspC(G457R) spores produce a CspC variant with an amino acid substitution of glycine 457 for arginine that render them hypersensitive to bile acid germinants and co-germinants. As a result, they exhibit increase spontaneous spore germination when plated on BHIS media lacking taurocholate germinant (24).

To measure the number of spores of each strain that spontaneously germinate in a sample, we pretreated spores with taurocholate or left them untreated and compared CFUs on media with and without taurocholate. Each strain was diluted to 2×10^7^ spores mL^-1^ (0.70 OD_600_ units in one mL) in a 1:1 mix of phosphate buffered saline and BHIS. Spores were then split into taurocholate-treated or untreated samples. The taurocholate-treated samples were incubated for 20 minutes at 37 °C in an aerobic environment to induce germination. This has previously been shown to not affect the CFUs recovered when plated on agar medium (23, 43). The pretreated and untreated spore samples were plated for a spore viability assay on both BHIS and BHIS+TA. To measure germination in GVA, all spores samples were diluted 10-fold and then embedded for GVA using both BHIS and BHIS+TA. With the concentrations of taurocholate used in the experiments, nearly all the spores will germinate synchronously in samples pre-treated with taurocholate and plated on or embedded in media without germinant relative to samples plated on or embedded in media with germinant (without prior exposure to bile acid germinant). Thus by comparing the CFU/mL calculated from these two different methods of inducing 100% germination, we will be able to determine if inducing germination before embedding in molten agarose media for GVA will decrease the viable spore count. Each of these assays was performed in biological triplicate.

Following overnight growth, colonies were counted, and the number of viable spores mL^-1^ were calculated (Fig. 4). GVA measurements were multiplied by 10 to account for the dilution performed. The general trends noting included 630Δ*erm* having higher germination rates when exposed to taurocholate, consistent with the current understanding of *C. difficile* germination. A slight increase was noted in Δ*cspC* when exposed to taurocholate. In ΔcspC/CspC(G57R), germination rates remain high even in the absence of taurocholate, but are slightly depressed as compared to exposure to taurocholate. For each condition, a t test compared the CFUs mL^-1^ measurements of GVA and the spore viability assay (Table 1). Across all conditions, no statistical difference was found. The relative germination was calculated for each condition by dividing the CFUs mL^-1^ measurement of the condition by either the CFUs mL^-1^ of the BHIS+TA measurement of either the same strain or 630Δ*erm*. No statistical difference between the coefficients of germination were found between GVA and the spore viability assay (Fig. 4). When 100% germination was set to the BHIS+TA pretreatment of the same strain, the average p value was 0.44 ± 0.29. When 100% germination was set to BHIS+TA pretreatment of 630Δerm was 0.55 ± 0.28.

**Figure 4:**
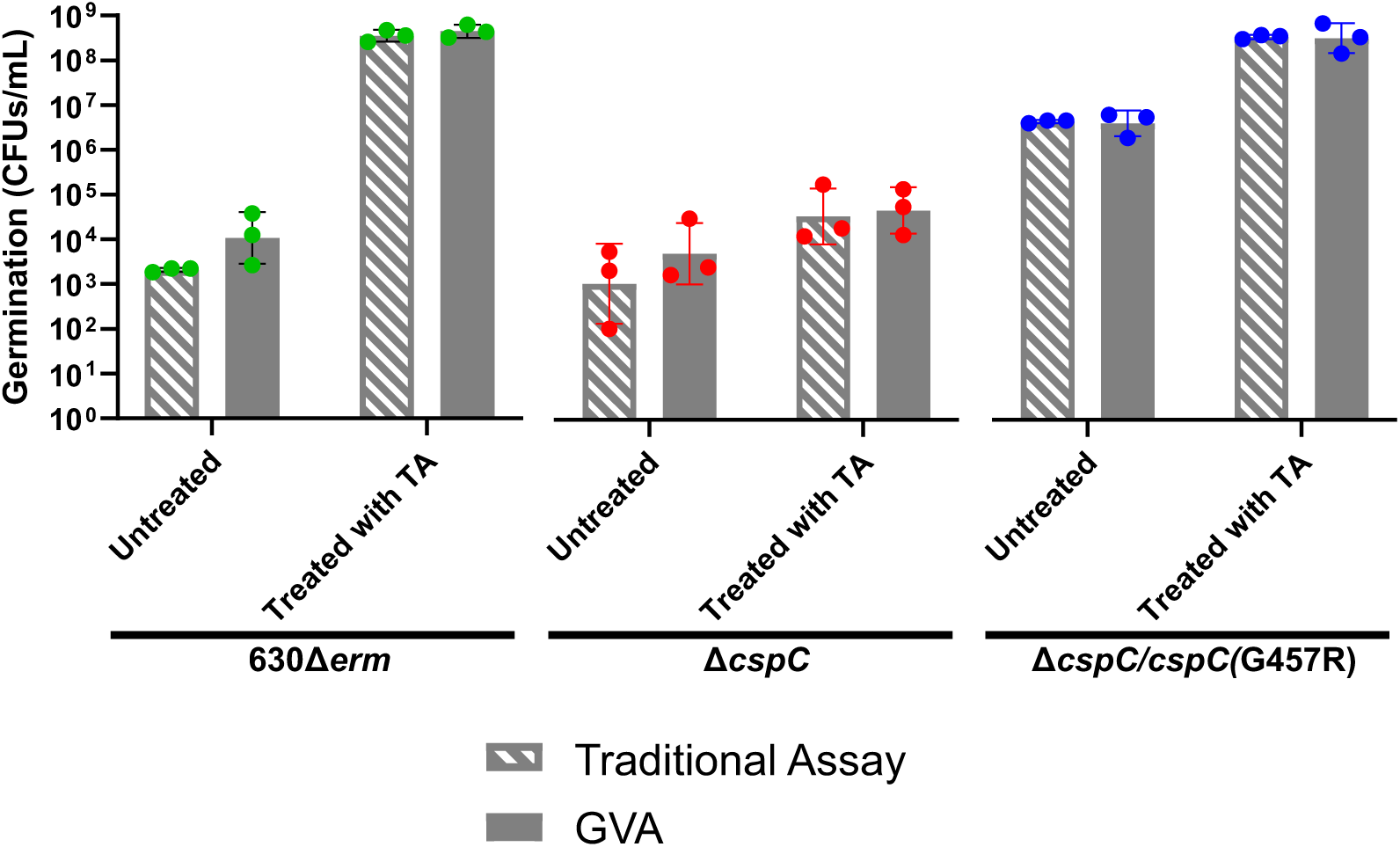
Measurements of spontaneous germination and full germination of three different strains of *C. difficile* were taken and compared. Spores isolated from indicated strains of *C. difficile* were pre-treated with either water or 1% taurocholate to induce germination and then plated (for traditional assay) or embedded in BHIS (for GVA) without germinant to enumerate spores that germinated during pre-treatment. Each bar is the average of three biological replicates, and error bars indicate the standard deviation for each condition.

**Figure 5:**
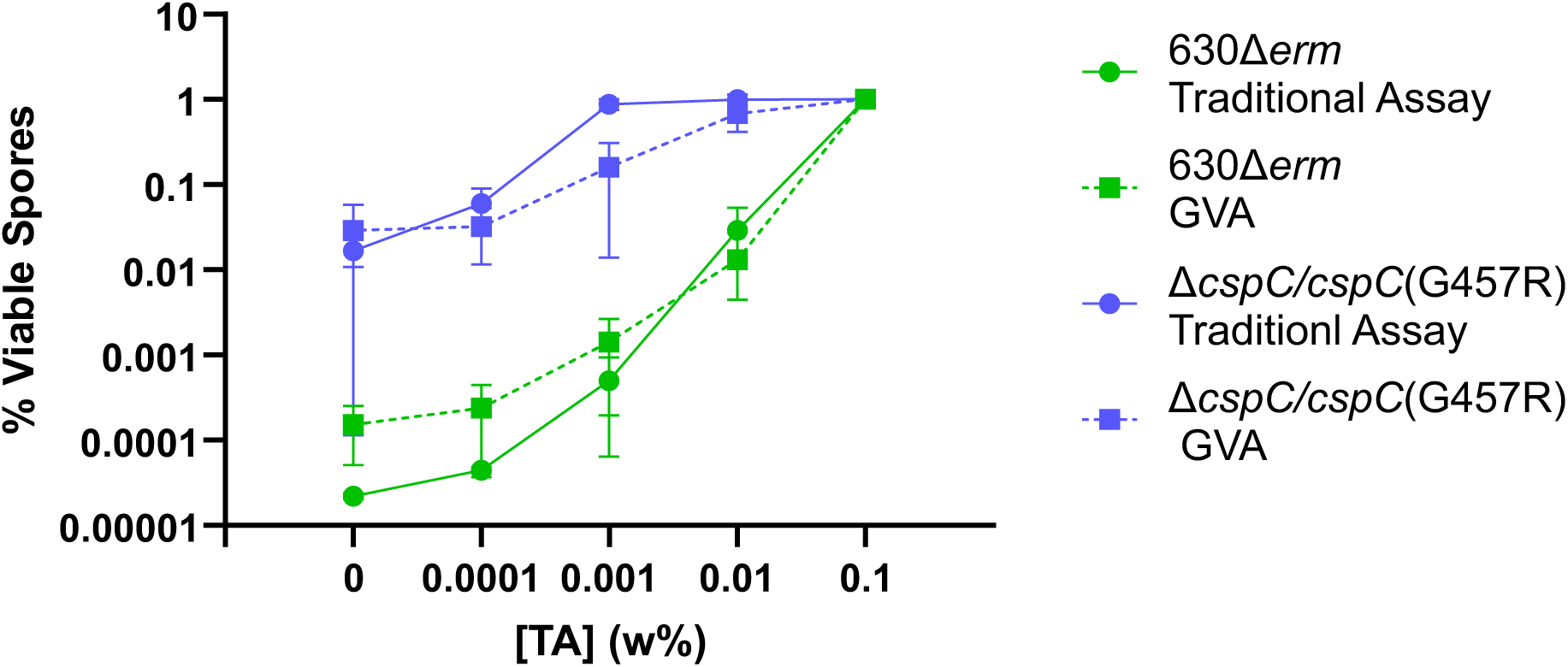
GVA measurements are similar to Spore Viability Assay measurements across multiple strains and germinant concentrations. Isolated wild-type spores (green) and spores with increased sensitivty to taurocholate (blue) were germinated at varying taurocholate concentrations ranging from 0.1% to 0.0001% and at 0% taurocholate. For the traditional assay (circles, solid line), spores were plated on BHIS media with the indicated taurocholate concentraion. For the GVA (squares, dashed line), spores were embedded in media with the indicated taurocholate concentration. Each point is the average of three biological replicates, and error bars show the standard deviations of the biological replicates at the given taurocholate concentration.

**Table 1:**
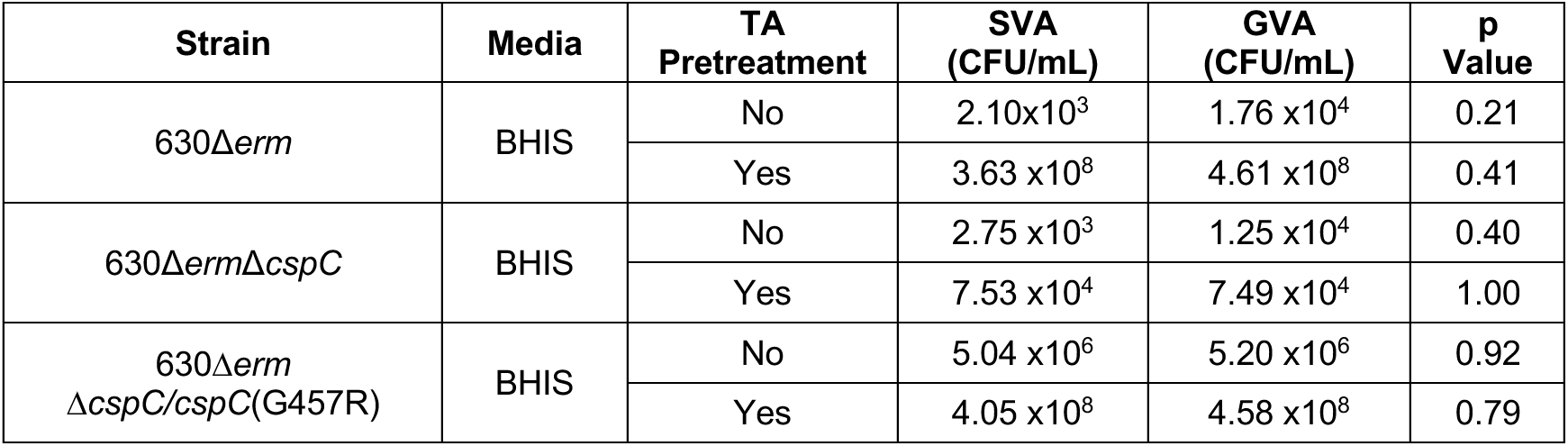
Both the taurocholate treated and untreated samples were diluted appropriately for both a GVA and spore viability assay. Each group was tested on both BHIS and BHIS+TA media. Following overnight growth, the spores/mL were calculated. A t-test was used to determine p values. The average across three biological replicates is presented above.

### GVA measures differences in strain sensitivity to germinant

Having shown that GVA can accurately measure the number of viable spores in a sample and the low levels of spontaneous germination that occur in rich media, we tested if the GVA could measure a difference in germinant sensitivity between strains. To test this, we compared the germination of 630Δ*erm* spores and Δ*cspC*/*CspC(G457R)* spores, which are hypersensitive to germinants, in the spot dilution spore viability assay and in GVA using BHIS medium containing various concentrations of taurocholate germinant. For each strain, 0.35 OD_600_ units of spores (approximately 10^7^ spores) were suspended in 1 mL of phosphate buffered saline. This sample was used for the spore viability assays and a 10-fold dilution of this sample was used for GVA. Each strain of *C. difficile* spores were plated on BHIS+TA plates containing increasing amounts of taurocholate concentrations of 0.0001% w/v (1.9 µM), 0.001% w/v (0.019 mM), 0.01% w/v (0.19 mM), 0.1% w/v (1.9 mM), or 0%, and allowed to grow overnight. The diluted sample was embedded in BHIS+TA medium containing the same concentrations of taurocholate for the GVA. For GVA, each concentration of taurocholate beyond 0.1% was formed by performing a 10-fold dilution of the previous concentration of taurocholate into fresh BHIS + agarose. Each assay was run in biological and technical triplicate.

After overnight growth, the colony counts of each assay were taken to calculate the germinated spores mL^-1^. The percentage of viable spores for each condition was calculated by dividing CFUs mL^-1^ measurement for a specific condition by the CFUs mL^-1^ measurement of the 0.1% w/v taurocholate condition for the corresponding strain and assay (Fig. 4). Visual inspection of the graph shows that the general trends between the two measurement types are similar across different strains. A paired t-test was used to determine statistical significance of the differences in the germination coefficients between GVA and spore viability assay measurements of the same strain at the same concentration of taurocholate. For all but one taurocholate concentration, no statistically significant differences were observed (p_min_=0.14, p_max_=0.61, p_mean_=0.28). The difference between the measurements of GVA and the spore viability assay were statistically significant for 630Δ*erm*Δ*cspC/cspC*(G457R) at a taurocholate concentration of 0.001% (p=0.017).

## Discussion

In this work, we assessed the utility of using GVA to measure the viability of and sensitivity to bile acid germinant of *C. difficile* spores. Since we found that the viable spore counts from GVA are highly comparable to the viable spore counts from the standard plate-based spore viability assay, our modified GVA is a promising alternative to the standard spore viability assay because it reduces the materials and time required to prepare each sample. Indeed, we further found that our modified assay detected spore CFUs with greater sensitivity. detecting spores 100-fold lower than the standard spore viability assay. GVA was also able to measure differences in germinant sensitivity between *C. difficile* strains and the low frequency of germination that occurs in the absence of germinants.

Overall, viable or germinated spore counts were quite similar between GVA and the traditional spore viability assay. The differences in germinated spore counts between assays, while not statistically significant, may be due to the differences between the two-dimensional nature of the spore viability assay and the three-dimensional nature of GVA. In GVA spores are embedded in the media with germinant and thus are exposed to germinant on all surfaces. In the spore viability assay, spores are plated on top of the agar media and thus are only exposed to germinate along one surface. If there are differences in germination rates due to this difference between two-dimensional versus three-dimensional exposure to germinates, we anticipate that germination rates measured by GVA would be more similar to what would occur in the biological context of the small intestine where spores would be exposed on all surfaces to germination signals.

In addition to measuring spore viability and germination in response to varying germinant concentrations, we also believe that GVA could be used to enumerate the number of vegetative *C. difficile* cells in a sample. However, our initial attempts to adapt GVA for use with vegetative *C. difficile* cells were only partially successful and led to some unexpected observations. At high CFUs mL^-1^, the GVA and CFU spot assay produced results with no statistical difference. However, at lower CFUs mL^-1^, no colonies grew in GVA and no growth was seen in BHIS liquid media even though viable cells were enumerated with the CFU spot assay. Such a large difference in the lower limit of detection between GVA and a CFU spot assay has not yet been reported in GVA literature (40, 41). It is possible these results point to a density-dependent growth of *C. difficile* in three-dimensional growth conditions, but further research is required to confirm this.

Recent research into the role and function of the Csp proteins during *C. difficile* germination has greatly increased our understanding of how this major nosocomial pathogen senses and responds to specific environmental signals to initiate infection (28, 29, 35, 44). However, questions remain about the specific mechanisms by which *C. difficile* spores sense germinant and co-germinant signals and propagate this signal to initiate germination. A clearer understanding of these processes could aid in the development of targeted therapies to specifically inhibit germination of *C. difficile* spores. Currently, research is limited by the lack of a high throughput method to measure germination and outgrowth of *C. difficile* spores. The current assay used to study these processes is a standard plate-based assay similar to a spot CFU assay, which has a high material cost and is time intensive, making high throughput assays difficult. The ability to adapt GVA to address different questions related to *C. difficile* spore germination could make this a powerful assay in further research into the mechanism of *C. difficile* spore germination and the development of specific inhibitors of *C. difficile* germination.

GVA is a highly adaptable assay that was able to be modified with relative ease to measure *C. difficile* spore viability and germination in a variety of conditions. The assay has also been shown to work in aerobic and anerobic conditions, with microbes ranging from bacteria to fungi to biofilms, and now with bacterial spores as well (40, 41). The simplicity of the technique enables researchers with basic laboratory skills, such as those typically taught in a general biology laboratory, to perform this assay with minimal additional training. Additionally, the low resource and time cost of the assay make it a potential candidate for adaptation for many other plate-based assays in microbiology. One such potential target is the plaque forming spot assay used to study bacteriophages. Fundamentally, the equations governing GVA could be used to replace any assay which uses serial dilutions. The most prominent limiting factor would be the microbes growing similarly in three-dimension as they do in standard two-dimensional conditions. While immediate applications of this technique are viability assay, the same principles could be applied to find concentrations of various particles in any solution. One potential adaptation would be a measurement of both live and dead cells in a solution. By staining cells with fluorescent markers, suspending them in an agarose matrix, and adapting a fluorescent microscope, one could map where in a subsection of the pipette tip’s volume the cells are found and utilize the GVA equations to determine the concentration of live, dead, and total cells within the solution. If this proposed technique works, then a slightly adapted version could also distinguish between different species. Overall, when GVA is broken down to the mathematics describing this assay, the principles become incredibly generalizable and may lead to more efficient assays and allow for new ways of measurements to be taken, allowing labs with access to different resources to make these measurements with less investment in new equipment.

## Methods

### Strains and Growth Conditions

*C. difficile* strain 630Δerm (23) was used unless otherwise noted. Strains 630ΔermΔcspC (23) and 630ΔermΔcspC/cspC(G457R) (24) were used during select experiments. All strains were gifts of the Shen Lab of Tufts University School of Medicine. Media plates for growth were either brain heart infusion supplemented with L-cystine (0.1% w/v; 8.25mM) and yeast extract (2.5 g per 500 mL) (BHIS) or on BHIS supplemented with taurocholate (0.1% w/v; 1.9mM)(BHIS+TA). Liquid cultures were grown in 3mL liquid BHIS without shaking. All bacterial growth occurred at 37°C under anerobic conditions using a gas mixture containing 85% N_2_, 5% CO_2_, and 10% H_2_. *C. difficile* spores isolated on different days were considered biological replicates. Technical replicates were considered multiple measurements from the same spore preparation.

### Sporulation

Sporulation was performed as previously described (24). Briefly, *C. difficile* inoculated from glycerol stocks was allowed to grow on BHIS+TA. The following day, one colony was inoculated into a 3mL BHIS liquid media overnight culture. The following morning, 50µL was back diluted into another 3mL BHIS liquid media culture and allowed to grow throughout the day. Sporulation was induced by spreading 120µL of the culture on 70:30 SMC:BHIS agar plates.

### Spore Purification

As previously described, sporulation was induced on 70:30 agar plates for 2-3 days(24). Following this, spores were collected and suspended in ice-cold water. Spore samples were washed with sterile water one time and allowed to sit at 4°C for 5 days. After this, the spores were washed with ice-cold water and then pelleted and treated with 20 units DNase I (RNase-free) (New England Biolabs). Spores were washed with ice-cold water again. Spores were then purified on a 20%/50% Histodenz gradient (Sigma Aldrich). The supernatant was removed, and the spores were washed three additional times before the pellet was resuspended in ice-cold water and stored at 4°C.

### SleC Cleavage Assay

SleC Cleavage Assay was performed as previously described (25). In brief, approximately 2 x 10^7^ spores (0.7 OD_600_ units) were diluted into 500 mL of water and 500 mL of liquid BHIS media was added. From this, 180µL were pipetted into a microcentrifuge tube containing 20µL of water for a negative control and 180µL were pipetted into a microcentrifuge tube containing 20µL 10% taurocholate (w/v). The microcentrifuge tube containing the taurocholate was incubated at 37°C for 20 minutes. Germination of solutions was quantified using either GVA or the spore viability assay. These assays were run with both BHIS and BHIS+TA media.

### Spore Viability Assay

Spore viability assay was performed as previously described (24, 32) with the following changes. Serial dilutions were prepared, either eight 1:10 dilutions or five 1:5 dilutions. 4.5µL droplets of each dilution were allowed to absorb onto the media. The plates were then inverted and placed into the incubator at 37°C. Plates were allowed to grow for 18-22 hours before plates were removed from the incubator and counted. All parts of this procedure were performed in the anerobic chamber, but the plates were removed for counting. Following counting, the number of colonies was multiplied by the dilution factor of its drop to calculate the CFUs in the drop. The value was then multiplied by 1000 and divided by 4.5 to calculate the CFUs/mL.

### GVA

#### Media Preparation

Medias for embedding were prepared according to standard protocols with the following changes. The amount of agarose added was changed to 0.66g per 100 mL for a concentration of 0.66% agarose. This allowed for a final agarose concentration of 0.5% (40). The media was then autoclaved on a liquid cycle. Following autoclaving, all non-autoclavable additives were added according to standard procedures. Media was allowed to solidify and stored at room temperature until use. Media can be microwaved three times and produce accurate results.

#### Preparation of samples

*C. difficile* spore samples were prepared in an aerobic environment. Approximately 1×10^7^ spores (0.35 OD_600_ units) were suspended in 1mL of PBS. This was then diluted 10-fold to a final concentration of 1×10^6^ spores mL^-1^. Any treatments to spores were made at this time. Then, 50 µL of treated sample were added a well of a 96 well plate.

#### Sample embedding

The sample embedding was performed as previously described with the following changes (40). TipOne pipette tips from USA Scientific were used for all trials. Prepared media was microwaved on high until completely liquid. It was then placed into a water bath at 55°C and allowed to reach thermal equilibrium. An aliquot of the media was poured into a reservoir and placed in a water bath inside of the anerobic chamber. 150 µL of media was mixed with 50 µL of sample and 150 µL of the resulting solution was carefully aspirated to avoid bubbles. The pipette tips were then placed into an ice bath for 18-20 seconds, after which they were placed into an empty pipette tip box. Pipettes were allowed to solidify at room temperature for 30 minutes before being placed into the incubator at 37°C overnight. Colony positions were measured the following day. Tips can be left at room temperature in an aerobic environment after growing overnight with no change in the imageability or the readings of the assay.

#### Tip imaging and colony counting

Imaging using a smart phone was performed as previously described (40). The 3D files for the optical configuration are available (www.printables.com/model/1709773-gva-modified-optical-setup). The phone stand was custom designed to fit with most phones and macro lens while the depth channel and depth bar are the designs of Meyer *et al* (40). Printing was done using a Formlabs Form4 resin printer with white v5 resin set to the default settings for this resin. Following printing, the resin was washed for 10 minutes in Form Wash (Formlabs) and dried for at least four hours. After drying, the pieces had a final cure of 30 minutes at 60°C in the Form Cure (Formlabs), the default time for this resin.

After printing, the holder was assembled, and the depth channel was secured to the stand base with a screw. A Samsung Galaxy S24 was fitted with a Xenvo macro lens with the wide-angle lens removed and placed into the phone holder of the optical configuration. The base of the configuration was positioned approximately 14.8 cm away from a white background. The tips were backlight with a Energizer motion sensing LED light. The images of the tips were then processed using the MATLAB app designed by Meyers et al (40).

## Bibliography

1. Abt MC, McKenney PT, Pamer EG. 2016. Clostridium difficile colitis: pathogenesis and host defence. Nat Rev Microbiol 14:609–620.

2. Centers for Disease Control and Prevention (U.S.). 2019. Antibiotic resistance threats in the United States, 2019. Centers for Disease Control and Prevention (U.S.).

3. Guh AY, Mu Y, Winston LG, Johnston H, Olson D, Farley MM, Wilson LE, Holzbauer SM, Phipps EC, Dumyati GK, Beldavs ZG, Kainer MA, Karlsson M, Gerding DN, McDonald LC. 2020. Trends in U.S. Burden of Clostridioides difficile Infection and Outcomes. N Engl J Med 382:1320–1330.

4. Desai K, Gupta SB, Dubberke ER, Prabhu VS, Browne C, Mast TC. 2016. Epidemiological and economic burden of Clostridium difficile in the United States: estimates from a modeling approach. BMC Infect Dis 16:303.

5. Magill SS, O’Leary E, Janelle SJ, Thompson DL, Dumyati G, Nadle J, Wilson LE, Kainer MA, Lynfield R, Greissman S, Ray SM, Beldavs Z, Gross C, Bamberg W, Sievers M, Concannon C, Buhr N, Warnke L, Maloney M, Ocampo V, Brooks J, Oyewumi T, Sharmin S, Richards K, Rainbow J, Samper M, Hancock EB, Leaptrot D, Scalise E, Badrun F, Phelps R, Edwards JR. 2018. Changes in Prevalence of Health Care–Associated Infections in U.S. Hospitals. N Engl J Med 379:1732–1744.

6. Theriot CM, Young VB. 2015. Interactions Between the Gastrointestinal Microbiome and *Clostridium difficile*. Annu Rev Microbiol 69:445–461.

7. Deakin LJ, Clare S, Fagan RP, Dawson LF, Pickard DJ, West MR, Wren BW, Fairweather NF, Dougan G, Lawley TD. 2012. The Clostridium difficile spo0A Gene Is a Persistence and Transmission Factor. Infection and Immunity 80:2704–2711.

8. Lim S-C, Hain-Saunders NMR, Imwattana K, Putsathit P, Collins DA, Riley TV. 2022. Genetically related Clostridium difficile from water sources and human CDI cases revealed by whole-genome sequencing. Environmental Microbiology 24:1221–1230.

9. Lim SC, Knight DR, Riley TV. 2020. *Clostridium difficile* and One Health. Clinical Microbiology and Infection 26:857–863.

10. Lawley TD, Clare S, Deakin LJ, Goulding D, Yen JL, Raisen C, Brandt C, Lovell J, Cooke F, Clark TG, Dougan G. 2010. Use of Purified Clostridium difficile Spores To Facilitate Evaluation of Health Care Disinfection Regimens. Applied and Environmental Microbiology 76:6895–6900.

11. Shen A. 2020. Clostridioides difficile Spores: Bile Acid Sensors and Trojan Horses of Transmission. Clin Colon Rectal Surg 33:058–066.

12. Baloh M, Sorg JA. 2022. *Clostridioides difficile* spore germination: initiation to DPA release. Current Opinion in Microbiology 65:101–107.

13. Kochan TJ, Foley MH, Shoshiev MS, Somers MJ, Carlson PE, Hanna PC. 2018. Updates to Clostridium difficile Spore Germination. Journal of Bacteriology 200:10.1128/jb.00218-18.

14. Shen A. 2020. *Clostridioides difficile* Spore Formation and Germination: New Insights and Opportunities for Intervention. Annu Rev Microbiol 74:545–566.

15. Lawler AJ, Lambert PA, Worthington T. 2020. A Revised Understanding of Clostridioides difficile Spore Germination. Trends in Microbiology 28:744–752.

16. Baloh M, Nerber HN, Sorg JA. 2022. Imaging Clostridioides difficile Spore Germination and Germination Proteins. J Bacteriol 204:e00210–22.

17. Dupuy B, Sonenshein AL. 1998. Regulated transcription of Clostridium difficile toxin genes. Molecular Microbiology 27:107–120.

18. Theriot CM, Koenigsknecht MJ, Carlson PE, Hatton GE, Nelson AM, Li B, Huffnagle GB, Z. Li J, Young VB. 2014. Antibiotic-induced shifts in the mouse gut microbiome and metabolome increase susceptibility to Clostridium difficile infection. Nat Commun 5:3114.

19. Francis MB, Allen CA, Shrestha R, Sorg JA. 2013. Bile Acid Recognition by the Clostridium difficile Germinant Receptor, CspC, Is Important for Establishing Infection. PLOS Pathogens 9:e1003356.

20. Sorg JA, Sonenshein AL. 2008. Bile Salts and Glycine as Cogerminants for Clostridium difficile Spores. Journal of Bacteriology 190:2505–2512.

21. Kochan TJ, Somers MJ, Kaiser AM, Shoshiev MS, Hagan AK, Hastie JL, Giordano NP, Smith AD, Schubert AM, Jr PEC, Hanna PC. 2017. Intestinal calcium and bile salts facilitate germination of Clostridium difficile spores. PLOS Pathogens 13:e1006443.

22. Galperin MY, Yutin N, Wolf YI, Alvarez RV, Koonin EV. 2022. Conservation and Evolution of the Sporulation Gene Set in Diverse Members of the Firmicutes | Journal of Bacteriology. Journal of Bacteriology. https://journals.asm.org/doi/10.1128/jb.00079-22. Retrieved 19 October 2026.

23. Kevorkian Y, Shen A. 2017. Revisiting the Role of Csp Family Proteins in Regulating Clostridium difficile Spore Germination. J Bacteriol 199.

24. Rohlfing AE, Eckenroth BE, Forster ER, Kevorkian Y, Donnelly ML, Benito De La Puebla H, Doublié S, Shen A. 2019. The CspC pseudoprotease regulates germination of Clostridioides difficile spores in response to multiple environmental signals. PLoS Genet 15:e1008224.

25. Kevorkian Y, Shirley DJ, Shen A. 2016. Regulation of Clostridium difficile spore germination by the CspA pseudoprotease domain. Biochimie 122:243–254.

26. Adams CM, Eckenroth BE, Putnam EE, Doublié S, Shen A. 2013. Structural and Functional Analysis of the CspB Protease Required for Clostridium Spore Germination. PLoS Pathog 9:e1003165.

27. Shen A, Edwards AN, Sarker MR, Paredes-Sabja D. 2019. Sporulation and Germination in Clostridial Pathogens. Microbiology Spectrum 7:GPP3-0017–2018.

28. McNellis ME, Pino GG-D, Serrano-Jiménez JA, Forster ER, Stoica AI, Heldwein EE, Shen A. 2026. The CspC:CspA heterodimer transduces germinant and co-germinant signals during Clostridioides difficile spore germination. PLOS Biology 24:e3003610.

29. Cal y Mayor Luna C, Alcorlo M, Kim C, Jacobson GN, Lázaro M, Valle M, Chang M, Hermoso JA, Mobashery S. 2026. Structure of core assembly of the Clostridioides difficile germinosome. Nat Commun 17:7669.

30. Gutelius D, Hokeness K, Logan SM, Reid CW. 2014. Functional analysis of SleC from Clostridium difficile: an essential lytic transglycosylase involved in spore germination. Microbiology 160:209–216.

31. Francis MB, Allen CA, Sorg JA. 2015. Spore Cortex Hydrolysis Precedes Dipicolinic Acid Release during Clostridium difficile Spore Germination. Journal of Bacteriology 197:2276–2283.

32. Fimlaid KA, Jensen O, Donnelly ML, Francis MB, Sorg JA, Shen A. 2015. Identification of a Novel Lipoprotein Regulator of Clostridium difficile Spore Germination. PLOS Pathogens 11:e1005239.

33. Paredes-Sabja D, Shen A, Sorg JA. 2014. Clostridium difficile spore biology: sporulation, germination, and spore structural proteins. Trends in Microbiology 22:406–416.

34. Donnelly ML, Fimlaid KA, Shen A. Characterization of Clostridium difficile Spores Lacking Either SpoVAC or Dipicolinic Acid Synthetase. https://journals.asm.org/doi/epub/10.1128/jb.00986-15. Retrieved 11 February 2026.

35. Bhattacharjee D, Francis MB, Ding X, McAllister KN, Shrestha R, Sorg JA. 2016. Reexamining the Germination Phenotypes of Several Clostridium difficile Strains Suggests Another Role for the CspC Germinant Receptor. Journal of Bacteriology 198:777–786.

36. Dembek M, Barquist L, Boinett CJ, Cain AK, Mayho M, Lawley TD, Fairweather NF, Fagan RP. 2015. High-Throughput Analysis of Gene Essentiality and Sporulation in Clostridium difficile. mBio 6:10.1128/mbio.02383-14.

37. Ribis JW, Melo L, Shrestha S, Giacalone D, Rodriguez EE, Shen A, Rohlfing A. 2022. Single-spore germination analyses reveal that calcium released during *Clostridioides difficile* germination functions in a feed-forward loop. preprint. Microbiology.

38. Wang S, Shen A, Setlow P, Li Y. 2015. Characterization of the Dynamic Germination of Individual Clostridium difficile Spores Using Raman Spectroscopy and Differential Interference Contrast Microscopy. J Bacteriol 197:2361–2373.

39. Donnelly ML, Forster ER, Rohlfing AE, Shen A. 2020. Differential effects of ‘resurrecting’ Csp pseudoproteases during Clostridioides difficile spore germination. Biochem J 477:1459–1478.

40. Meyer CT, Lynch GK, Stamo DF, Miller EJ, Chatterjee A, Kralj JM. 2023. A high-throughput and low-waste viability assay for microbes. Nat Microbiol 8:2304–2314.

41. El-Fateh M, Meyer CT, Chatterjee A, Zhao X. 2025. A high-throughput anaerobic method for viability assays. Microbiology Spectrum 13:e02706–24.

42. Meyer CT, Lynch GK, Stamo DF, Miller EJ, Chatterjee A, Kralj JM. 2023. A high-throughput and low-waste viability assay for microbes. Nat Microbiol 8:2304–2314.

43. Dembek M, Stabler RA, Witney AA, Wren BW, Fairweather NF. 2013. Transcriptional Analysis of Temporal Gene Expression in Germinating Clostridium difficile 630 Endospores. PLoS ONE 8:e64011.

44. Roseiro I, Nunes A, Martins D, Alves F, Persson S, Henriques AO, Oleastro M, Serrano M. 2026. A Single Amino Acid Substitution in CspA Increases Germination Sensitivity and Broadens Bile Salt Germinant Specificity in Clostridioides difficile Spores. Environmental Microbiology 28:e70332.

